# Monetary feedback modulates performance and electrophysiological indices of belief updating in reward learning

**DOI:** 10.1101/183665

**Authors:** Daniel Bennett, Karen Sasmita, Ryan T. Maloney, Carsten Murawski, Stefan Bode

## Abstract

Belief updating entails the incorporation of new information about the environment into internal models of the world. Bayesian inference is the statistically optimal strategy for performing belief updating in the presence of uncertainty. An important open question is whether the use of cognitive strategies that implement Bayesian inference is dependent upon motivational state and, if so, how this is reflected in electrophysiological signatures of belief updating in the brain. Here we recorded the electroencephalogram of participants performing a simple reward learning task with both monetary and non-monetary instructive feedback conditions. Our aim was to distinguish the influence of the rewarding properties of feedback on belief updating from the information content of the feedback itself. A Bayesian updating model allowed us to quantify different aspects of belief updating across trials, including the size of belief updates and the uncertainty of beliefs. Faster learning rates were observed in the monetary feedback condition compared to the instructive feedback condition, while belief updates were generally larger, and belief uncertainty smaller, with monetary compared to instructive feedback. Larger amplitudes in the monetary feedback condition were found for three event-related potential components: the P3a, the feedback-related negativity (FRN) and the late positive potential (LPP). These findings suggest that motivational state influences inference strategies in reward learning, and this is reflected in the electrophysiological correlates of belief updating.

## 1. Introduction

In learning and decision making, finite cognitive resources should be allocated so as to optimise decisions about behaviourally relevant outcomes, while minimising expenditure of resources on irrelevant or inconsequential tasks (Boureau, Sokol-Hessner, & Daw, 2015; Collins & Frank, 2012; Pitz & Sachs, 1984; Simon, 1976; Stenhav et al., 2017). One cognitive domain in which this trade-off arises is belief updating, the process by which relevant new information is integrated with prior beliefs to produce a new belief state that can be used to formulate appropriate responses to events in one’s environment (Achtziger, Alós-Ferrer, Hügelschäfer, & Steinhauser, 2014; Gläscher, Daw, Dayan, & O’Doherty, 2010; Wunderlich, Dayan, & Dolan, 2012). In environments characterised by uncertain contingencies or unpredictable dynamics, belief updating has obvious survival implications; however, many of the cognitive and neurophysiological factors arbitrating this component of decision-making remain unclear. In particular, one open question is how belief updating might be affected by the presence of reward, and whether reward directly impacts on motivational state and, by doing so, improves belief updating (Achtziger, Alós-Ferrer, Hügelschäfer, & Steinhauser, 2015; Correa et al., 2018). This question is of particular importance given the ongoing debate in educational psychology and behavioural economics regarding the efficacy of using rewards to incentivise performance (Hidi, 2016; Lazear, 2000).

Bayesian theories of cognition propose that the cognitive processes underlying human decision making in a variety of tasks can be accurately explained as a form of Bayesian inference (e.g., Austerweil, Gershman, Tenenbaum, & Griffiths, 2015; Chater & Oaksford, 2008; Friston et al., 2015; Knill & Pouget, 2004; Körding, 2007; Ma & Jazayeri, 2014; Richards & Knill, 1996). Bayesian inference is the statistically optimal strategy for unbiased belief updating (Bennett, Murawski, & Bode, 2015; Kolossa, Kopp, & Fingscheidt, 2015; Nassar, Wilson, Heasly, & Gold, 2010; Stern, Gonzalez, Welsh, & Taylor, 2010). For a behaving agent, Bayesian belief updating involves repeatedly revising a prior belief distribution based on incoming sensory data, where at each timepoint belief about the probability of an event (posterior distribution) is proportional to the product of the agent’s previous belief (prior distribution) and the likelihood of the event given sensory data. Using a Bayesian modelling framework, Bennett et al. (2015) recently developed a novel reward learning paradigm that required participants to use information provided by stimulus feedback to update their beliefs after each trial. In particular, participants in this task learned the association between the contrast levels of a dynamically changing checkerboard stimulus and monetary reward values, which provided an indication of the location of a hidden ‘target’ contrast. A simple Bayesian model provided dynamic estimates of the belief states of each participant on each trial, which allowed for trial-by-trial quantification of (a) the uncertainty of beliefs, and (b) the magnitude of belief updates. Importantly, these estimates of belief state were found to correlate with the amplitude of components of the event-related potential (ERP), including a positive relationship between P3 amplitudes and belief update magnitude that was not accounted for by simpler models (Bennett et al., 2015). This finding was consistent with earlier theories and other empirical findings linking the P3 and Bayesian belief updating (Kolossa et al., 2015; Kopp, 2008).

While Bayesian models often fit learning behaviour well overall, their goodness-of-fit can deteriorate sharply as the cognitive demands of Bayesian inference increase (Payzan-LeNestour & Bossaerts, 2011), or when simpler, and less effortful, heuristic strategies conflict with Bayesian inference (Achtziger et al., 2015; Charness & Levin, 2005). This suggests that discrepancies between Bayesian models and human behaviour may be in part motivational: since full Bayesian inference can be computationally demanding, participants’ motivation to engage in effortful cognition may moderate their use of strategies that resemble Bayesian belief updating (Lieder & Griffiths, 2015; Shenhav, Botvinick, & Cohen, 2013). Since an important factor that directly modulates motivation is the availability of reward, this motivational account predicts that the use of monetary performance incentives in learning tasks (as in the study by Bennett et al. (2015)) may affect the behavioural strategies employed by participants. Monetary incentive has been linked with improved performance in various learning paradigms (e.g., Bonner & Sprinkle, 2002; Kleih, Nijboer, Halder, & Kübler, 2010), but the effect of monetary incentives on participants’ use of Bayesian belief updating is an open question. In addition, we would predict that any motivational influences on belief updating should also manifest in the electrophysiological indices of belief updating, such as the P3 ERP component. In preliminary support of this notion, P3 amplitudes are also known to be influenced by the motivational value of stimuli (Sato et al., 2005).

In the present study, we investigated the effect of motivational state, manipulated in terms of financial reward value, in a reward learning task with graded feedback similar to that introduced by Bennett et al. (2015). Here, however, our approach allowed us to disambiguate belief updating based on the information content provided at feedback itself, from the reward value associated with that feedback. In this task, feedback was delivered in the form of either monetary reward (monetary feedback condition; as in Bennett et al., 2015) or simple instructional directives, where no extrinsic reward was provided (the instructive feedback condition). Importantly, feedback values were constrained such that the information content of feedback was identical across the two feedback conditions. As in Bennett et al. (2015), we employed a Bayesian modelling framework to quantify trial-by-trial aspects of belief updating, namely belief update magnitude and belief uncertainty.

Further, we probed the neural mechanisms underlying belief updating in the two feedback conditions by examining three ERP components associated with motivation, learning and/or reward processing. We note that our aim was not to conduct an exhaustive exploration of the role of Bayesian inference in reward learning, which is an active topic of research in the machine learning literature (e.g., Ghavamzadeh, Mannor, Pineau & Tamar, 2015). Rather, we aimed to examine the influence of certain higher-level motivational factors (tied to reward availability) on human belief updating, and to measure the potential electrophysiological correlates of this influence. For this, we investigated three ERP components: the P3 (Bennett et al., 2015; Goldstein et al., 2006; Jepma et al., 2016; Kleih et al., 2010; Polich, 2007; Sato et al., 2005), the feedback related negativity (FRN; Yeung & Sanfey, 2004), and the late positive potential (LPP; Ito, Larsen, Smith, & Cacioppo, 1998). The connection between individuals’ revision of probabilistic beliefs and the amplitude of the P3 is consistent with the hypothesis that it reflects a Bayesian belief-updating mechanism (Bennett et al., 2015; Kolossa et al., 2015; Kopp, 2008; Mars et al., 2008). This theory suggests that P3 amplitudes reflect the degree to which internal models of the environment are updated, and predicts that larger P3 amplitudes would be tied to greater engagement of cognitive strategies that implement Bayesian belief updating. Since the increase in P3 amplitude has been shown to be proportional to the gradual increase of monetary feedback (Goldstein et al., 2006; Kleih et al., 2010; Sato et al., 2005), we hypothesised that higher P3 amplitudes following monetary feedback would be observed in addition to an increase elicited by the amount of belief updating (the update magnitude). Traditionally, the P3 has been partitioned into the frontocentral P3a subcomponent, and the centroparietal P3b subcomponent (Polich, 2007). Specifically, it is the P3a that has been associated with the magnitude of belief updating (Bennett et al., 2015). This suggestion is also consistent with a functional imaging study indicating the frontal encoding of belief update size at the anterior cingulate cortex (O’Reilly et al., 2013), a region considered to be a potential substrate for the P3a (Polich, 2007). The Bayesian account of P3a amplitude is generally consistent with the prominent context-updating theory of the P3, which argues that P3 amplitude is an index of the revision of schemata concerning stimulus context (Donchin & Coles, 1988). This revision process is analogous to the updating of prior beliefs in the Bayesian sense. FRN amplitude was also investigated because of its importance as an index of outcome evaluation in reward learning and feedback processing (Achtziger et al., 2015; Frank, Woroch, & Curran, 2005; Holroyd & Coles, 2002). Finally, in research studying the processing of emotional stimuli, LPP amplitude is thought to differentially encode positive and neutrally-valenced stimuli (Keil et al., 2002; Schupp et al., 2000); we therefore sought to investigate whether LPP amplitude differed between monetary and instructive feedback conditions.

## 2. Method and Materials

### 2.1 Participants

Twenty-three participants were recruited from among students of the University of Melbourne, Australia (mean age = 23.40; age range 19-31; 17 female, six male). Participants were right-handed, had normal or corrected-to-normal visual acuity and no medical history of any neurological disorder. Informed consent was acquired from all participants in accordance with the Declaration of Helsinki, and approval was obtained from the University of Melbourne Human Research Ethics Committee (ID 1339694). Participants received monetary compensation for participation, and in addition they received a financial reward that was proportional to task winnings in the monetary feedback condition only (mean = $25.24; SD = 4.05, for details, see below). For all participants, total remuneration value was within the range AUD $20-30.

Four participants were excluded from analysis of EEG data: one because of an excessive number of artefacts (more than 80 percent of trials affected), one because of a failure of the eyeblink artefact removal routine, and two additional participants because of computer error during EEG acquisition. Final EEG analyses were therefore performed on data provided by 19 participants (mean age = 23.75; age range 19-31; 13 female, six male).

### 2.2 Behavioural task

Participants performed a reward learning task modified from that first introduced by Bennett et al. (2015) while EEG was recorded. This task required participants to learn the fixed target contrast of a dynamically changing greyscale checkerboard stimulus (Figure 1A) on the basis of feedback presented across subsequent trials. On each trial, the checkerboard was presented for up to 30 seconds during which time its contrast changed linearly (alternately increasing and decreasing, changing direction at upper/lower contrast bounds).

**Figure 1.**
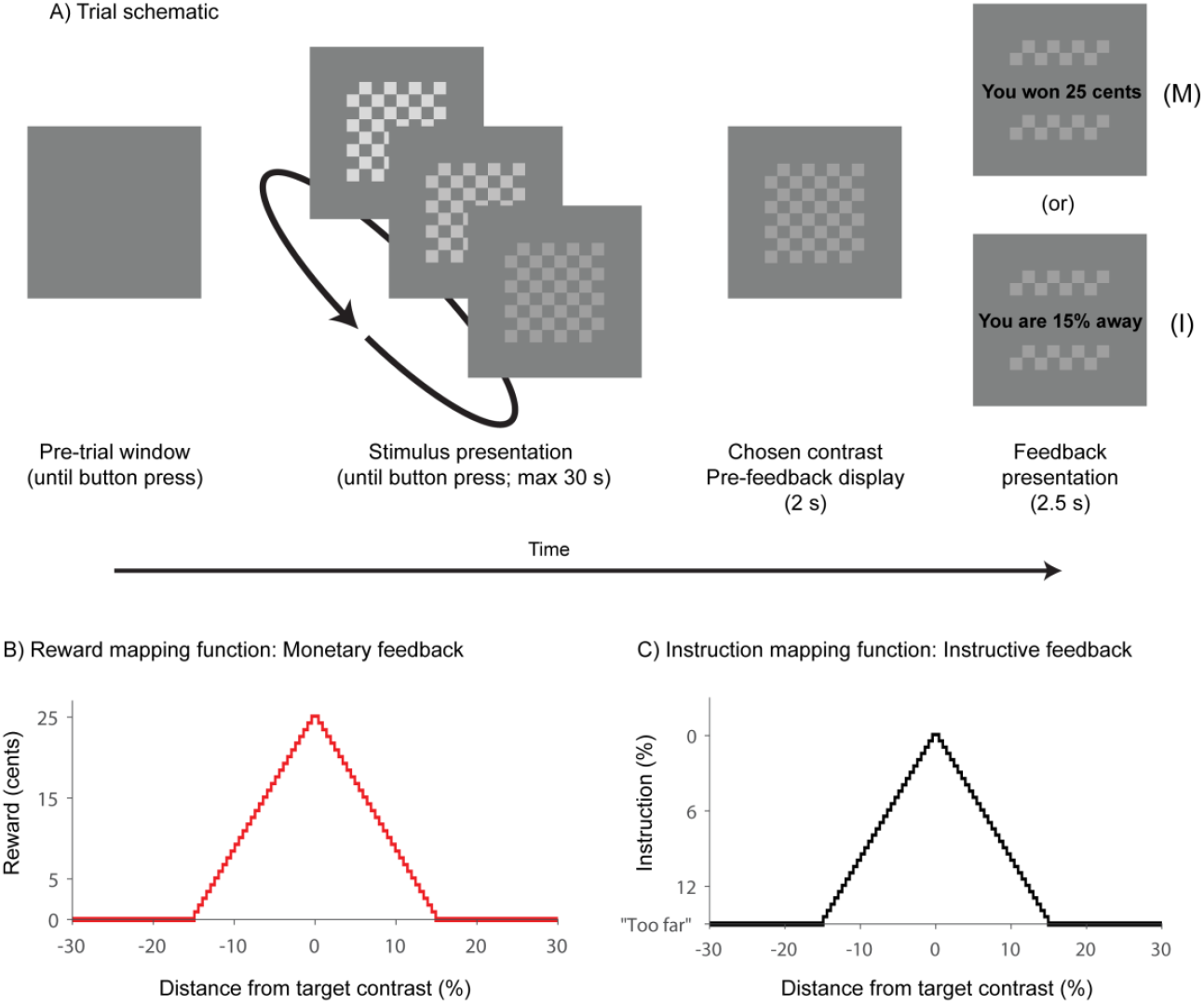
(A) Trial schematic. Following a self-paced button press, a checkerboard stimulus was presented with a linearly changing contrast. The participant could at any time select the contrast displayed on screen by pressing a button with the right index finger. The trial continued until a button was pressed, or until stimulus duration exceeded 30 seconds. Following the participant’s choice, the selected contrast remained on screen for two seconds, after which time the monetary (M) or instructive (I) feedback associated with the chosen contrast was displayed for 2.5 seconds. In the event that no button was pressed within 30 seconds, feedback was a message reminding the participant of the task instructions. (B) Feedback mapping for monetary feedback condition. The mapping was a symmetrical triangular function with a centre of zero percent contrast difference, a half-width of 15 percent contrast difference, and a height of 25 cents. As such, received reward was maximal when the participant responded at the target contrast, and decreased with increasing difference of chosen contrast from the target. Reward was zero for responses at greater than 15 percent distance. Feedback received was rounded to the nearest whole-cent value. (C) Feedback mapping for the instructive feedback condition. For responses at less than 15 percent difference from the target, participants were informed of the difference between the chosen contrast and the target contrast (rounded to the nearest of 49 equally spaced values, in order to match precisely the step size of the monetary condition’s feedback mapping). For responses at greater than 15 percent difference from the target, participants were informed only that their response was “too far” from the target (equivalent to the monetary condition’s zero cent feedback).

At the start of a block of trials, participants were initially unaware of the target contrast. By a process of trial and error, they selected a contrast on each trial and were given visual feedback on how close their choice was to the hidden target contrast. On each trial, initial contrast, initial direction of contrast change (increasing/decreasing), and rate of change were randomised using the same parameters as in Bennett et al. (2015). At any time during stimulus presentation, the participant could choose the contrast displayed on screen by pressing a button with the right index finger. After a two second delay in which the chosen contrast remained on screen, participants received feedback regarding their chosen contrast. The present study employed a novel variant of this task in which feedback regarding the target contrast could be either monetary (as in the original paradigm of Bennett et al., 2015) or purely instructive. In the monetary condition, this feedback was presented in the form of monetary reward (e.g., “You won 15 cents”) according to a triangular function *M* (Equation 1) of the distance between the chosen and the target contrast (see Figure 1B):

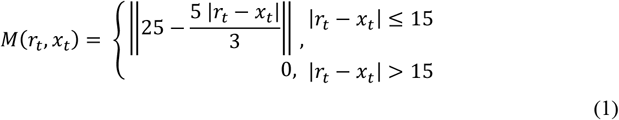

where *t* is the trial number in the sequence, *rt* is the target contrast on trial *t*, and *x_t_* is the participant’s chosen contrast on trial *t*. Double bars denote rounding to the nearest integer cent. Responses closer to the target contrast earned proportionally more (up to a maximum of 25 cents per trial, rounded to the nearest integer cent), while participants received zero reward for responses at greater than 15 percent contrast distance from the target.

In the instructive condition, feedback took the form of an explicit instructional directive informing the participant of the distance between their chosen contrast and the target (e.g., “You were 11.25% away from the target”; see Figure 1C), with no monetary incentive attached. For responses at greater than 15 percent distance from the target contrast, participants were informed only that their response was “too far” from the target. As such, in the instructive feedback condition the reward mapping function *M* from Equation 1 was replaced with the instruction mapping function *I*:

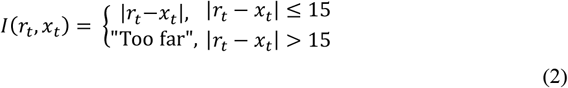

Crucially, to ensure strict equivalence in the feedback information provided between the instructive and monetary feedback conditions, instructive feedback values were constrained to follow an equivalent functional form to monetary feedback (compare Figures 1B and 1C). This was done by rounding instructive feedback values to the nearest value in the set {0, 0.625, 1.25, 1.875,…, 15}. For any given sequence of choices, therefore, feedback in the two conditions provided identical information regarding the target contrast. Consequently, any differences in task performance between instructive and monetary conditions cannot be attributed to differences in the information content of feedback.

Participants completed a total of 14 blocks of trials of the task over approximately 50 minutes. Monetary and instructive conditions were presented in seven consecutive blocks each, with the starting condition counterbalanced across participants. Each block had a different target contrast, selected pseudo-randomly from the interval [25%, 85%]. Blocks ran until cumulative checkerboard presentation duration exceeded three minutes, or until 25 trials were completed, whichever occurred sooner. As a result, the number of trials per block varied across participants, ensuring that they could not hurry through the task in an attempt to trade off experiment duration against monetary winnings. Upon receiving feedback, participants were not informed of the exact numerical contrast level of their choice; instead, the checkerboard remained on-screen at the chosen contrast while written feedback was presented (Figure 1A). As a result, learning was necessarily affected by perceptual uncertainty regarding the identity of the chosen contrast. Prior to the task, participants were trained in interpretation of feedback in both feedback conditions, and testing commenced only when a satisfactory level of task understanding was evident.

Stimuli were presented using a Sony Trinitron G420 CRT monitor at a framerate of 120 Hz. During task performance, participants were seated comfortably in a darkened room, using a chin rest at a fixation distance of 77 cm from the screen. Checkerboard stimuli were 560 × 560 pixels in size, measuring 19.5 × 19.5 cm on the screen and subtending14.43 × 14.43 degrees of visual angle. Responses were recorded using a five-button Cedrus Response Box. All other task parameters were identical to those employed by Bennett et al. (2015), with the exception that the checkerboard in the present task did not phase-reverse, and therefore had a smoothly changing (rather than flickering) appearance.

### 2.3 Model-based estimates of belief updating

A Bayesian grid estimator (Moravec, 1988), as described and implemented for the reward learning task used by Bennett et al. (2015), was used to quantify dynamic aspects of belief updating in terms of the degree of participants’ belief uncertainty on a given trial, and the magnitude of belief updating across trials as new feedback information was imparted. This estimator calculated a probabilistic estimate of participants’ beliefs regarding the level of the target contrast in each trial and used this belief distribution to estimate choice likelihoods. Formally, beliefs were described by a probability mass function *θ* over a contrast space discretised into *J* equally sized bins, where the value of the function *θ* at each bin represented the participant’s subjective probability that the target contrast (denoted *rt*) fell within bin *j* on trial *t*. Bins had a width of 0.61% contrast, resulting in a belief distribution that contained *J* = 148 contrast bins on the interval [10, 100]. This value was chosen because it was the largest value sufficient to resolve different values of monetary feedback. On each trial *t*, participants observed the feedback *f_t_* after the choice of contrast bin *x_t_*, determined according to the monetary and instructive feedback mapping functions *M* and *I* as specified by Equations 1 and 2, respectively. Belief estimates were initialised in each block as a discrete uniform distribution, representing participants’ *a priori* uncertainty regarding the target contrast level. This belief distribution was then updated sequentially according to Bayes’ Rule as feedback was received, such that the posterior distribution of trial *t* formed the prior distribution for trial *t* + 1:

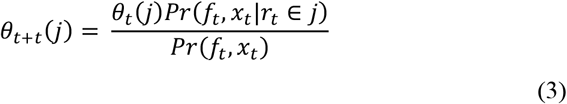

The left-hand side of Equation 3 represents the posterior belief distribution for contrast bin *j* following trial *t*, and is calculated by multiplying the participant’s prior belief that the target contrast fell within bin *j, θ_t_*(*j*) by the likelihood of observing the choice/feedback pair (*f_t_, x_t_*) if the target were in bin *j, Pr*(*f_t_, x_t_|r_t_* ∈ *j*), and dividing by the marginal likelihood of the update, *Pr*(*f_t_, x_t_*).

As noted above, variability in task performance between participants was captured by the response uncertainty parameter *σ*. Formally, *σ* represents the standard deviation of the Gaussian noise affecting belief updates after feedback receipt, such that larger values of *σ* indicate a greater degree of noise in the updating process, and therefore more imprecise belief updates. Since participants were not informed of the numerical value of the contrast they had chosen but had to estimate this chosen contrast from the visual display, this response uncertainty therefore also results in a Gaussian prior over chosen contrast. For a complete discussion of the mathematical role of *σ* in the Bayesian updating model, see Bennett et al. (2015).

To estimate choice likelihood, this model used a probability of maximum utility choice rule (cf. Speekenbrink & Konstantinidis, 2015), whereby contrast bins with a higher probability of containing the target contrast had a proportionally higher probability of being chosen, subject to response uncertainty during choice:

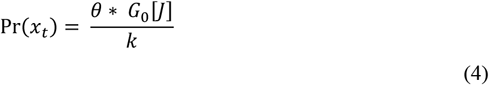

As such, on each trial the choice likelihood probability mass function was determined by convolving the prior belief distribution by the uncertainty function *G*_0_ over the set of contrast bins *J*, where *k* is a normalisation constant and square parentheses denote the domain of convolution. Intuitively, this response model implies that response probabilities are derived by the addition of Gaussian noise to the target contrast distribution *J*. The uncertainty function G0 was a zero-mean Gaussian function of the contrast difference between the true chosen contrast *x_t_* and each bin *x_j_* of the distribution *θ*. This function was also parameterised by *σ* (truncated to the available range of contrasts):

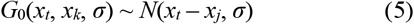

In the implementation of the Bayesian grid estimator used in the present study, the likelihood function reflected the participant’s uncertainty regarding which contrast he or she had chosen. The implication of this is that if a participant had perfect knowledge of exactly which contrast he or she had chosen, *Pr*(*f_t_,x_t_*) would simply be chosen such that the posterior distribution integrated to unity. However, here we instead made the more plausible assumption that participants had imperfect knowledge of which contrast they chose (for example, if the true chosen contrast was 50 percent, the participant might ‘know’ only that they had chosen some contrast between 40 and 60 percent). Under these circumstances, Bayesian principles dictate that the participant ought to weight the belief update by their uncertainty regarding the chosen contrast. This can be achieved by considering the marginal likelihood not as a point value but as a set of candidate contrasts varying in subjective probability and integrating across this set to determine the magnitude of the belief update. In the present study, we modeled the response uncertainty as Gaussian noise around the true chosen contrast, with mean zero and standard deviation determined as a parameter *σ* fit individually to participants. The parameter *σ* was estimated with maximum likelihood estimation using the MATLAB Optimization Toolbox (The Mathworks, Natick, MA), and was fit separately to instructive and monetary conditions for each participant. In estimating *σ* we assumed that the Gaussian noise distribution was truncated such that zero probability was assigned to contrast bins that were not displayed during the experiment (i.e. those outside the interval [10, 100]).

Model estimations of subjective belief distributions were used to calculate two variables of interest on each trial: belief uncertainty prior to the receipt of feedback, and post-feedback belief update magnitude. These allowed us to represent participants’ belief states as a function of previously observed evidence, and the subjective information content of feedback. A formal encapsulation of the model can be found in Bennett et al. (2015). Briefly, we defined belief uncertainty as the Shannon entropy (Shannon, 1948) over contrast bins of the prior distribution:

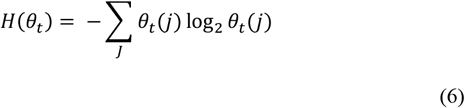

Because the entropy *H* of a probability distribution reflects the uncertainty coded by those probabilities, higher values of entropy in the belief distribution correspond to greater levels of belief uncertainty. An entropy of zero is observed only in the case of complete certainty, where all probabilities in the distribution but one are zero. The entropy in a distribution is maximal when all probabilities are equal, as in a uniform distribution (Bennett et al., 2015).

Belief update magnitude was calculated as the mutual information of feedback. This quantity provides an indication of the degree by which uncertainty has been resolved in the updating from prior (before feedback) to posterior (after feedback) probabilities. It corresponds to the information content (*V*) of the feedback: more informative feedback promotes a greater reduction in uncertainty from prior to posterior beliefs. Larger values of *V* indicated a greater resolution of uncertainty (and hence, a larger belief update). We calculated mutual information as the difference in entropy between prior and posterior belief distributions:

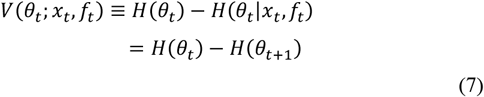

Both belief uncertainty and belief update magnitude thus provided model-based estimates that allowed us to compare the updating of beliefs across trials in the two feedback conditions. Namely, an estimate of the reduction in uncertainty of beliefs as a result of feedback (belief uncertainty), and the degree to which that feedback was usefully applied by participants in updating their beliefs about the location of the target contrast in contrast space (belief update magnitude). We anticipated that feedback in the form of monetary reward would yield larger reductions in belief uncertainty, as well as belief updates of greater magnitude, when compared to instructive feedback. Multiple linear regression was used to examine whether belief uncertainty and trial number (an indicator of learning improvement with time) could predict behavioural performance in terms of the choice error (the percent contrast difference between the chosen and target contrasts). Regressions were performed across trials for each participant separately, and the resulting *β* coefficients were subjected to single-sample *t*-tests to assess whether the effect of the predictors was significantly greater than zero across participants.

### 2.4. EEG data acquisition

The electroencephalogram was recorded from 35 Ag/AgCl active scalp electrodes (Fp2, AF7, AF3, AFz, AF4, AF8, F5, F1, Fz, F2, F4, F6, FC1, FCz, FC4, FC6, C5, C3, Cz, C4, CP5, CP3, CP1, CPz, CP6, P5, P1, Pz, P4, P6, POz, PO8, O1, Oz, Iz in the International 10-20 System). Electrodes interfaced with a BioSemi ActiveTwo 64-channel system running ActiView acquisition software and used an implicit reference during recording. Due to technical problems with electrode hardware, not all 64 channels could be recorded for all participants. Therefore, based on previous (Bennett et al., 2015) and planned analyses, data was acquired from prespecified channels of interest, including all frontocentral and centroparietal midline electrodes. All electrode channels included in subsequent event related potential (ERP) analyses were recorded without issue for all participants, and data quality was not compromised. Data were linearly detrended and re-referenced offline to an average of left and right mastoid electrodes. The vertical and horizontal electrooculogram (EOG) were recorded from electrodes infraorbital and horizontally adjacent to the left eye. EEG was recorded at a sampling rate of 512 Hz.

Preprocessing of data was performed using a semi-automated preprocessing pipeline (cf. Bode, Bennett, Stahl, & Murawski, 2014; Brydevall, Bennett, Murawski, & Bode, 2018). Data were first manually screened to exclude epochs contaminated by skin potential or muscle artefacts. Using a linear FIR filter, data were then highpass filtered at 0.1 Hz, lowpass filtered at 70 Hz, and notch filtered at 50 Hz to remove background electrical noise. Epochs were generated consisting of data from 1500 milliseconds before to 1500 milliseconds after feedback presentation. An independent components analysis (ICA), as implemented in the EEGLAB toolbox (Delorme & Makeig, 2004) for MATLAB, was performed on the resulting dataset to identify and remove components related to eye movements and eyeblink artefacts. Finally, an automatic artefact screening procedure excluded all epochs from analysis in which maximum/minimum amplitudes exceeded ±200 *μ*V.

### 2.5 ERP data analysis

We assessed three ERP components: the P3a, the FRN, and the LPP. Component amplitudes were calculated using estimation routines implemented in the ERPlab plugin (Lopez-Calderon & Luck, 2014), time-locked to feedback presentation on each trial and baseline-corrected from 0 to 500 ms pre-feedback. P3a amplitude was calculated as the largest positive peak in the window from 250-550ms post-feedback at the frontocentral and centroparietal midline electrodes AFz, Fz, FCz, Cz, and CPz (Bennett et al., 2015). This time window allowed us to estimate peak amplitude within a symmetrical window about the peak of the P3a as identified in grand average waveforms.

At the same midline electrodes, FRN amplitude was calculated as the peak-to-peak distance between the most negative peak in the window from 200 to 550 ms and the immediately preceding positive peak (Achtziger et al., 2015; Frank et al., 2005; Yeung & Sanfey, 2004). A peak-to-peak measure of the FRN was used rather than a mean amplitude measure to ensure that estimates of FRN and P3a amplitude were statistically independent of one another. Finally, LPP amplitude was calculated as the mean voltage within the window from 550 to 900 ms post-feedback at the centroparietal midline electrodes Cz, CPz, and Pz (Hajcak, Dunning, & Foti, 2009; Ito et al., 1998). This time window was chosen both to accord with previous literature (e.g., Keil et al., 2002), and to ensure that P3a and LPP analysis windows did not overlap.

The effects of motivational factors on ERP signatures of belief updating were assessed in three-way repeated-measures ANOVAs (2×2×5 for the P3a and FRN, 2×2×3 for the LPP) with fixed within-group factors of feedback condition (monetary and instructive), belief update magnitude (small and large: determined according to a median split over trials; see below) and electrode scalp location (five levels for P3a and FRN analyses: AFz, Fz, FCz, Cz, and CPz; three levels for LPP analyses: Cz, CPz, and Pz), while participant served as a random factor.

## 3. Results

### 3.1 Behavioural results

Participants completed a variable number of trials per block (mean = 17.57; SD = 2.70). Paired-samples *t*-tests found no evidence to suggest that the number of trials completed per block differed between instructive and monetary feedback conditions, *t*(22) = .80, *p* = .43, and no evidence to suggest that the total stimulus viewing time differed between feedback conditions, *t*(22) = −1.04, *p* = .31. These analyses suggest that participants adopted an equivalent speed/accuracy trade-off in both feedback conditions. Behavioural performance was quantified by choice error, defined as the absolute difference between the chosen contrast and the target contrast on each trial. We investigated differences in choice error as a function of trial number and feedback condition using linear mixed-effects analysis with feedback condition and trial number as fixed effects. Results indicated a significant main effect of trial number, *F*(24,60.95) = 13.09, *p* < 0.0001, with performance generally improving over time within each block, indicating acceptable overall task performance (see Figure 2A). There was also a significant main effect of feedback condition, *F*(1,9.29) = 20.07, *p* = 0.001, driven by better overall learning performance in the monetary than the instructive feedback condition. We found no evidence for block-order effects for either the monetary (*t*(21) = 1.48, *p* = .15) or the instructive feedback condition (*t*(21) = −1.13, *p* = .27). Finally, the interaction between feedback condition and trial number was also significant, *F*(24, 60.95) = 2.89, *p* = 0.0004. This effect is likely to have been driven by greater differences between monetary and instructive feedback conditions in mid-block trials, rather than in block-initial trials (Figure 2A). Such a pattern stands to reason, since participants began each block with no *a priori* knowledge regarding the target contrast and were just as likely to make a correct as an incorrect initial guess regardless of feedback condition. In addition, given the demand characteristics of the task being performed by participants, it is to be anticipated that the later stages of each block would be associated with performance approaching an asymptotic level (i.e., not declining fully to 0 error). The level of this asymptote is likely to depend upon multiple factors, including limits on performance associated with perceptual uncertainty (related to participants’ limited ability to distinguish between nearby contrast values), and working memory limitations (related to the limited precision with which a given contrast value can be stored in working memory during learning).

**Figure 2.**
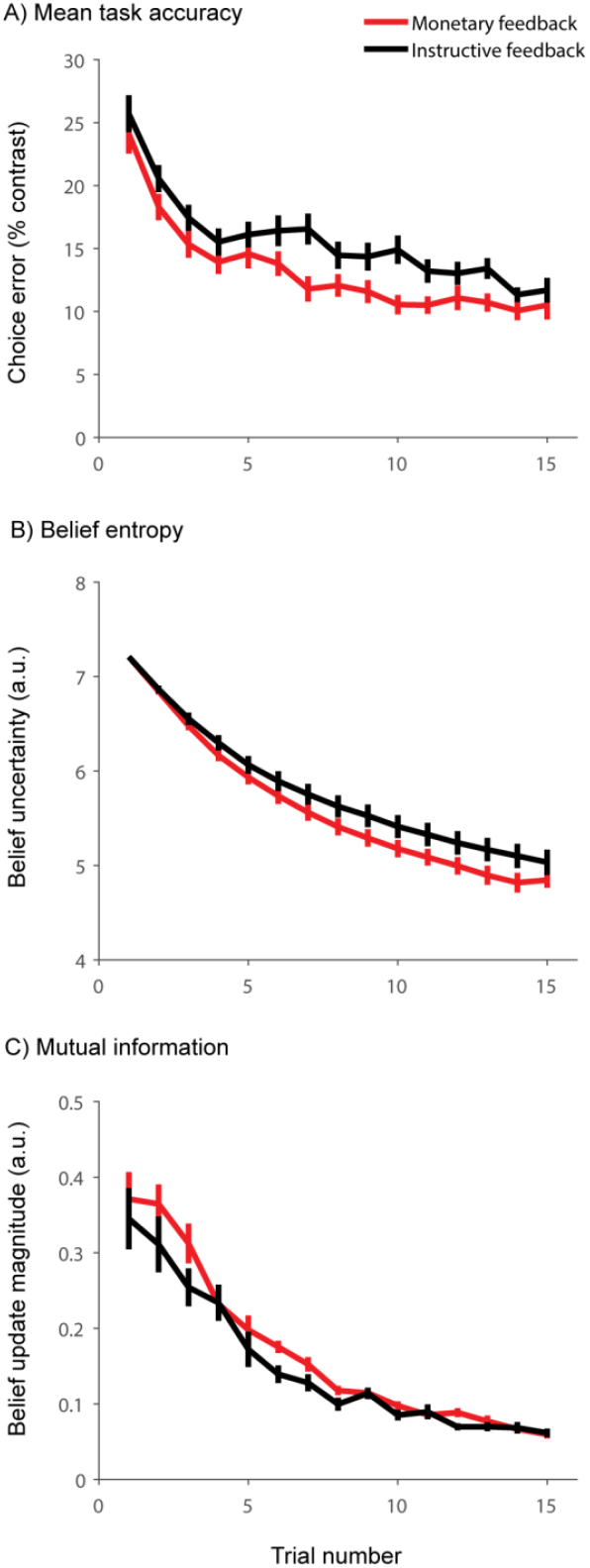
Overall reward learning task performance and model-derived belief variables as a function of trial number across participants (*n* = 23). In all plots, monetary feedback is displayed in red, instructive feedback is displayed in black. (A) Mean choice error (measured as absolute difference between chosen and target contrasts) as a function of feedback condition and trial number. (B) Belief entropy, giving an estimate of belief uncertainty. (C) Mutual information, giving an estimate of belief update magnitude. Note that, in all plots, since data for block-wise averages were not available for all participants for trials 16 – 25, only trials 1-15 are displayed here. Error bars represent the standard error of the mean.

### 3.2 Modelling results

Figures 2B and 2C display the mean model estimates for pretrial belief uncertainty (Shannon entropy, *H*, see Equation 6) and belief update magnitude (mutual information, *V*, see Equation 7) as a function of trial number, respectively. Multiple linear regression was used to predict the behavioural task accuracy, or choice error (the absolute difference between the chosen and target contrasts) on the basis of trial number and belief uncertainty. With monetary feedback, both trial number, mean *β* = −0.004, *t*(21) = −6.21, *p* < 0.0001, and belief uncertainty, mean *β* = 0.04, *t*(21) = 8.61, *p* < 0.0001, were significantly related to the choice error. Similarly, with instructive feedback, *β* coefficients for trial number, mean *β* = −0.005, *t*(21) = −7.59, *p* < 0.0001 and belief uncertainty, mean *β* = 0.05, *t*(21) = 8.96, *p* < 0.0001, significantly differed from zero, indicating that on average they were significantly related to behavioural choice error on the task. Overall, these analyses indicated that belief uncertainty, as estimated from the Bayesian inference model, was related to participants’ performance, even when trial-by-trial effects of learning (as indicated by trial number) were accounted for in the regression analysis. In other words, on trials with greater belief uncertainty, there tended to be a larger error in participants’ choice of stimulus contrast. This provides validation for the use of the Bayesian model-derived estimates of belief variables in subsequent analyses.

In the Bayesian updating model, participants’ response uncertainty was captured by the parameter *σ*, the standard deviation of the Gaussian noise affecting the marginal likelihood of belief updates. Estimates of *σ* across participants had a mean value of 13.4 (SD = 4.0) in the monetary feedback condition and 17.22 (SD = 12.43) in the instructive feedback condition. A paired-samples *t*-test found that this difference was not statistically significant, *t*(21) = 1.51, *p* = .15, perhaps due to the relatively high variance in estimates of *σ* in the instructive feedback condition. Values in the monetary feedback condition closely mirrored those obtained by Bennett et al. (2015) in their version of the task. As identified by Bennett et al. (2015), response uncertainty estimates were positively correlated with participants’ overall task performance, as measured by the mean absolute difference between the chosen and target contrasts across trials for both the monetary feedback condition, *r*(20) = 0.81, *p* < 0.0001, and the instructive feedback condition, *r*(20) = 0.64, *p* = 0.001. In both conditions therefore, on average individuals with less response uncertainty (smaller values of *σ*) tended to choose contrasts closer to the target contrast, further validating our choice of the Bayesian grid estimator in describing participants’ beliefs during the task.

### 3.3 ERP results

We next investigated whether any of the three relevant ERP components displayed analogous patterns to those observed in the behavioural and modelling results (see Figure 2). We also took the measures of belief update magnitude as estimated by the model and, by way of a median split, divided trials into either “large” or “small” update magnitude categories. This analysis allowed us to identify electrophysiological indices which were associated with the differential relative performance between monetary and instructive feedback, as well as any that might reflect the updating of belief states across trials.

Scalp maps for the P3a and LPP analysis windows are presented in Figure 3 and depict the grand average voltage topography across participants for the monetary and instructive feedback conditions, as well as their difference. Figure 3G shows the average waveform across participants at a representative electrode (Fz) for both feedback conditions. Analyses of peak latencies of each ERP component indicated no significant effects of feedback condition or belief update magnitude, at any electrode assessed. A cursory inspection of Figure 3 suggests that, in general, mean voltage tended to be of a higher amplitude for the monetary feedback condition compared to the instructive feedback condition in both the P3 (250-550 ms) and LPP (550-900 ms) analysis windows. Mean amplitudes for each ERP component according to feedback condition, belief update magnitude and electrode site are given in Figure 4.

**Figure 3.**
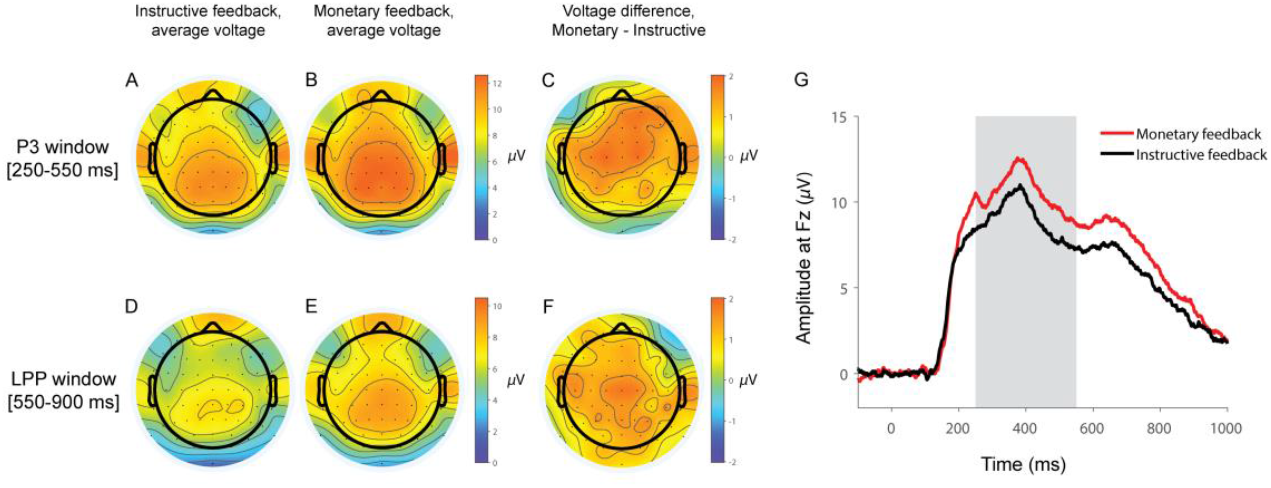
Scalp maps of grand average voltages for monetary and instructive feedback conditions as well as their difference (in *μ*V), for the different analysis windows, as well as the mean waveform for the two feedback conditions at a representative electrode (*n* = 19). Top row (A-C): P3 analysis window (250-550 ms post-feedback onset). Bottom row (D-F): LPP analysis window (550-900 ms post-feedback onset). Left column (A and D): Mean voltage in the instructive feedback condition. Centre column (B and E): Mean voltage in the monetary feedback condition. Right column (C and F): Mean voltage differences across the two conditions. For all scalp maps, voltages at missing electrodes have been reconstructed using spline interpolation for display purposes only. (G): Grand average feedback-locked ERP waveforms at electrode Fz, grouped by feedback condition (red: monetary feedback; black: instructive feedback). Time 0 denotes the onset of task performance feedback. The grey shaded region highlights the P3 analysis window used here.

**Figure 4.**
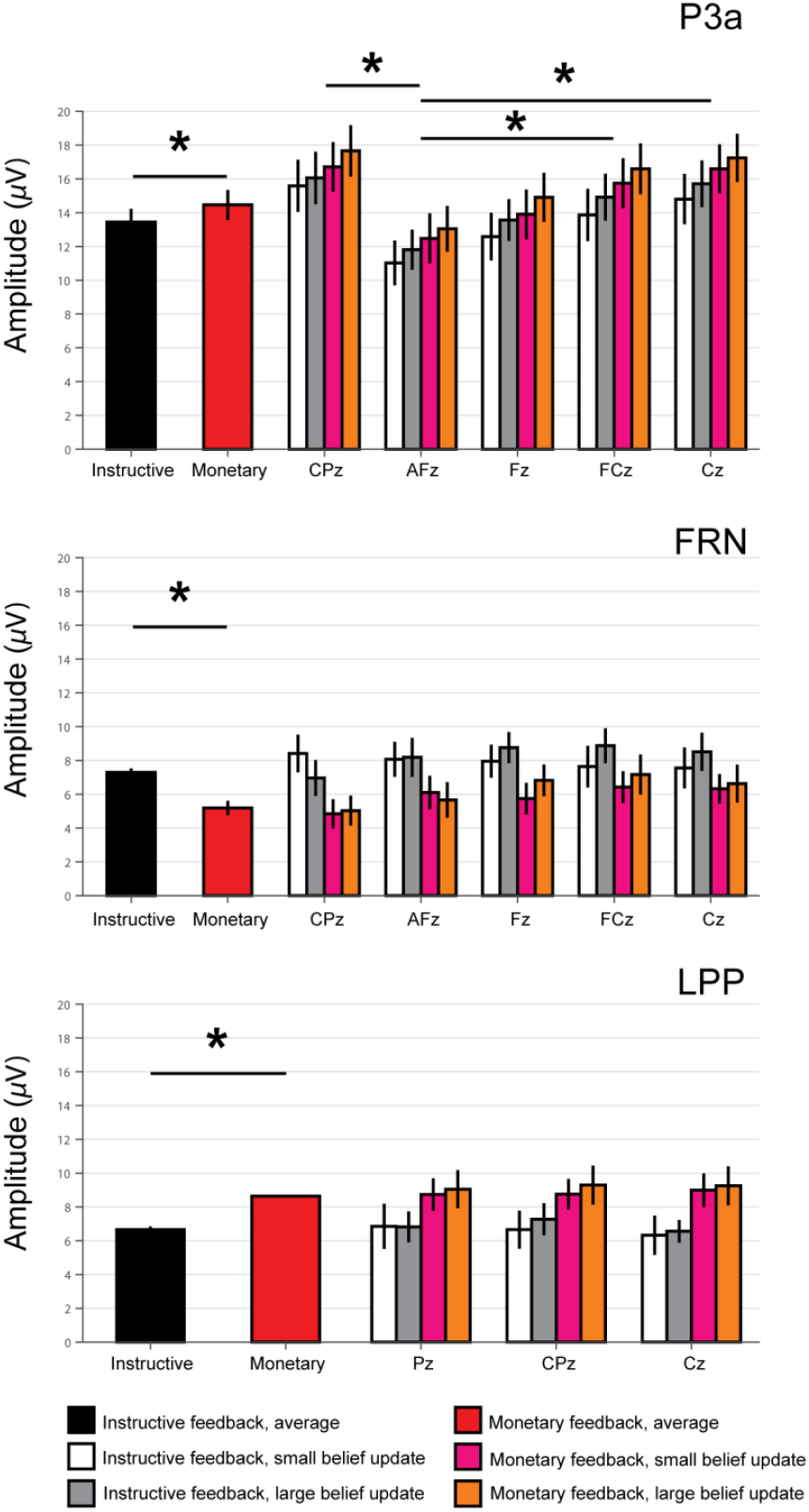
Mean ERP component amplitudes as a function of feedback condition, electrode site and belief update magnitude (divided into two bins by way of a median split: large or small, according to model-derived estimates). The two wide bars on the left show mean amplitudes marginalised across electrodes (black bars: instructive feedback; red bars: monetary feedback). Grouped bars to the right show the same data separately for electrode site and belief update magnitude (white bars: instructive feedback, small update; grey bars: instructive feedback, large update; pink bars: monetary feedback, small update; orange bars: monetary feedback, large update). In all plots, error bars represent the standard error of the mean. * indicates *p* < 0.05. (A) P3a amplitudes at electrodes CPz, AFz, Fz, FCz, and Cz. We observed a significant main effect of feedback condition, such that monetary feedback was associated with larger P3a components than instructive feedback, in addition to a main effect of electrode site. (B) FRN amplitude at electrodes CPz, AFz, Fz, FCz, and Cz. FRNs were significantly larger in the instructive feedback condition than the monetary feedback condition. (C) LPP amplitude at electrodes Pz, CPz, and Cz. We observed a significant effect of feedback condition, such that monetary feedback was associated with larger LPP components than instructive feedback.

#### 3.3.1 P3a

A 2×2×5 repeated-measures ANOVA with factors of feedback condition (instructive, monetary), belief update magnitude (small, large), and electrode site (AFz, Fz, FCz, Cz, CPz) revealed a significant main effect of feedback condition, *F*(1,360) = 5.44, *p* = 0.02, 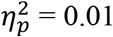, and of electrode site, *F*(4,360) = 6.43, *p* = 0.0001, 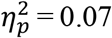, but not of belief update magnitude, *F*(1,360) = 1.63, *p* = 0.20, on mean P3a peak amplitude. The main effect of feedback condition was driven by the overall greater P3a amplitudes for the monetary compared to the instructive conditions (see Figure 4A). For the main effect of electrode site, Bonferroni-corrected pairwise comparisons indicated that the mean P3a amplitudes at CPz, FCz and Cz were all significantly greater than that at AFz (*p* = 0.0002, *p* = 0.02 and *p* = 0.001, respectively), across feedback conditions and belief update magnitudes. Although the grouped bars of Figure 4A suggests that there was a trend for an interaction between feedback condition and belief update magnitude, with P3a amplitudes being generally larger with larger belief updates, and larger still with monetary feedback, the interaction term was not significant, *F*(1,360) = 0.0001, *p* = 0.98. Similarly, the other two-way interaction and the three-way interaction were all non-significant (*p* > 0.10).

#### 3.3.2 FRN

A 2×2×5 repeated-measures ANOVA with within-groups factors of feedback condition (instructive, monetary), belief update magnitude (small, large), and electrode (AFz, Fz, FCz, Cz, CPz) indicated a significant main effect of feedback condition on FRN amplitude, *F*(1,360) = 18.91, *p* < 0.0001, 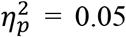, with larger FRNs elicited by instructive than rewarding feedback (see Figure 4B). Since in the present study the FRN was measured as a peak-to-peak change in voltage, this finding indicates that the FRN component decreased further from the preceding positive peak (i.e. became more negative) in the instructive condition than in the rewarding condition. All other main effects and interactions were non-significant (all *p* > 0. 10).

#### 3.3.3 LPP

A 2×2×3 repeated measures ANOVA with within-groups factors of feedback condition (instructive, monetary), belief update magnitude (small, large), and electrode (CPz, Pz, Cz) revealed a significant main effect of feedback condition on mean LPP amplitude, *F*(1,216) = 13.99, *p* = 0.0002, 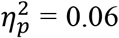. As with the P3a peak amplitudes, this effect denotes the larger overall mean amplitude for the monetary feedback condition compared to the instructive feedback condition (Figure 4C). There were no significant main effects of belief update magnitude, *F*(1,216) = 0.28, *p* = 0.60, or electrode site, *F*(2,216) = 0.28, *p* = 0.96, and none of the two-way interactions nor the three-way interaction were significant (all *p* > 0. 10).

### 3.4 Regression analyses

Although the Bayesian model-derived estimates of belief update magnitude did not influence mean amplitudes of any of the ERP components investigated, it is possible that model-derived belief state variables could serve as a predictor for neural signatures of the underlying updating process when the actual value of monetary reward is accounted for. We therefore conducted multiple linear regressions for each ERP component, electrode site, and feedback condition, to test for whether model-derived belief state variables (belief update magnitude and belief uncertainty) might serve as predictors for ERP amplitudes when the value of reward is also encapsulated in the statistical model. Regressions were performed separately for monetary and instructive feedback conditions, using reward value (fixed at zero in the instructive condition) and model-derived belief update magnitude as predictors for P3a and LPP amplitudes. Because the FRN has been linked to the evaluation of feedback outcomes and the magnitude of a reward prediction error associated with reward learning (Achtziger et al., 2015; Holroyd & Coles, 2002; Miltner, Braun, & Coles, 1997; Yeung & Sanfey, 2004), we expected it to reflect belief uncertainty rather than belief update magnitude; hence we tested its utility as a predictor of FRN amplitude, along with reward value. In addition, trial number was included as an additional predictor in all regression analyses to ensure that any observed relationships were not the result of time-on-task effects within each block.

*R*^2^ and *β* coefficients derived from each regression are given in Tables 1–3 for each ERP component below. In short, for monetary feedback, belief update magnitude was positively related to FRN amplitude at the central midline electrode sites CPz and Cz when reward value, trial number, and belief uncertainty were accounted for. For the LPP, belief uncertainty was positively associated with amplitude at electrode Cz, and reward value was significantly related to LPP amplitude at all electrodes tested (Pz, CPz, and Cz). Belief update magnitude, however, was not significantly related to P3a amplitude, and trial number was not significantly related to any ERP component at any electrode. There were no significant regression models for instructive feedback. This, however, was expected since in those, reward value was fixed at zero and thus could not account for any of the variance (results not reported here). In addition, the pattern of regression results was not substantively altered if Kullback-Leibler divergence was used instead of mutual information as a measure of belief update magnitude (cf. O’Reilly et al., 2013).

**Table 1:** Regression analyses of P3a component in Monetary feedback condition

|  | CPz |  |  |  | AFz |  |  |  | Fz |  |  |  | FCz |  |  |  | Cz |  |  |  |
| --- | --- | --- | --- | --- | --- | --- | --- | --- | --- | --- | --- | --- | --- | --- | --- | --- | --- | --- | --- | --- |
| | $\beta$ | $t$ | $p$ | $R^2$ | $\beta$ | $t$ | $p$ | $R^2$ | $\beta$ | $t$ | $p$ | $R^2$ | $\beta$ | $t$ | $p$ | $R^2$ | $\beta$ | $t$ | $p$ | $R^2$ |
| Reward value | 150.89 | 1.34 | 0.20 | 0.18 | 139.19 | 1.34 | 0.20 | 0.17 | 158.21 | 1.49 | 0.16 | 0.19 | 182.69 | 1.71 | 0.11 | 0.25 | 163.69 | 1.67 | 0.12 | 0.27 |
| Belief update magnitude | 45.48 | 0.89 | 0.39 |  | 34.17 | 0.73 | 0.48 |  | 30.6 | 0.64 | 0.53 |  | 44.96 | 0.93 | 0.37 |  | 57.12 | 1.29 | 0.22 |  |
| Belief uncertainty | 8.13 | 1.09 | 0.29 |  | 7.66 | 1.12 | 0.28 |  | 7.18 | 1.02 | 0.32 |  | 9.87 | 1.40 | 0.18 |  | 10.35 | 1.60 | 0.13 |  |
| Trial number | -0.11 | -0.41 | 0.69 |  | -0.19 | -0.76 | 0.46 |  | -0.18 | -0.70 | 0.50 |  | -0.24 | -0.90 | 0.38 |  | -0.21 | -0.91 | 0.38 |  |

$R^2$ values are for the regression model as a whole, including the intercept term.

**Table 2:**
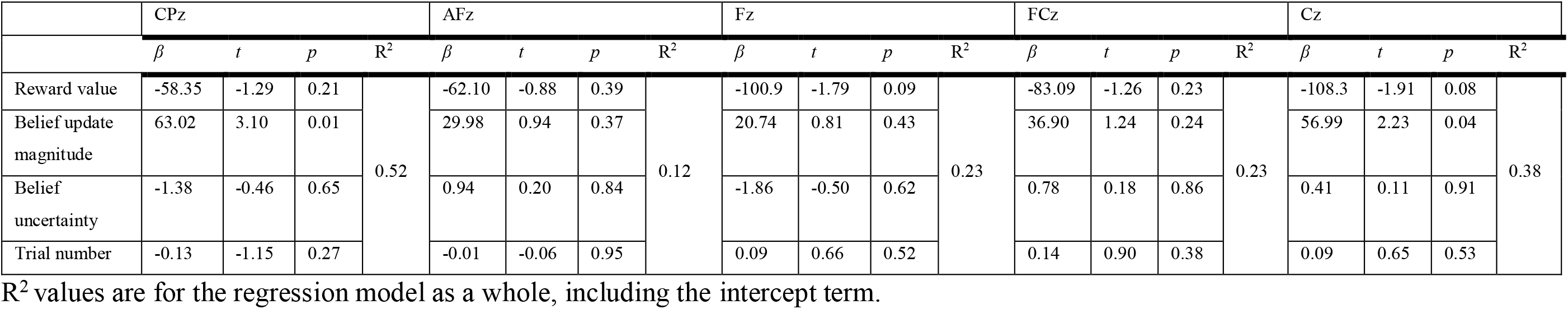
Regression analyses of FRN component in Monetary feedback condition

|  | CPz |  |  |  | AFz |  |  |  | Fz |  |  |  | FCz |  |  |  | Cz |  |  |  |
| --- | --- | --- | --- | --- | --- | --- | --- | --- | --- | --- | --- | --- | --- | --- | --- | --- | --- | --- | --- | --- |
| | $\beta$ | $t$ | $p$ | $R^2$ | $\beta$ | $t$ | $p$ | $R^2$ | $\beta$ | $t$ | $p$ | $R^2$ | $\beta$ | $t$ | $p$ | $R^2$ | $\beta$ | $t$ | $p$ | $R^2$ |
| Reward value | -58.35 | -1.29 | 0.21 | 0.52 | -62.10 | -0.88 | 0.39 | 0.12 | -100.9 | -1.79 | 0.09 | 0.23 | -83.09 | -1.26 | 0.23 | 0.23 | -108.3 | -1.91 | 0.08 | 0.38 |
| Belief update magnitude | 63.02 | 3.10 | 0.01 |  | 29.98 | 0.94 | 0.37 |  | 20.74 | 0.81 | 0.43 |  | 36.90 | 1.24 | 0.24 |  | 56.99 | 2.23 | 0.04 |  |
| Belief uncertainty | -1.38 | -0.46 | 0.65 |  | 0.94 | 0.20 | 0.84 |  | -1.86 | -0.50 | 0.62 |  | 0.78 | 0.18 | 0.86 |  | 0.41 | 0.11 | 0.91 |  |
| Trial number | -0.13 | -1.15 | 0.27 |  | -0.01 | -0.06 | 0.95 |  | 0.09 | 0.66 | 0.52 |  | 0.14 | 0.90 | 0.38 |  | 0.09 | 0.65 | 0.53 |  |

$R^2$ values are for the regression model as a whole, including the intercept term.

**Table 3:**
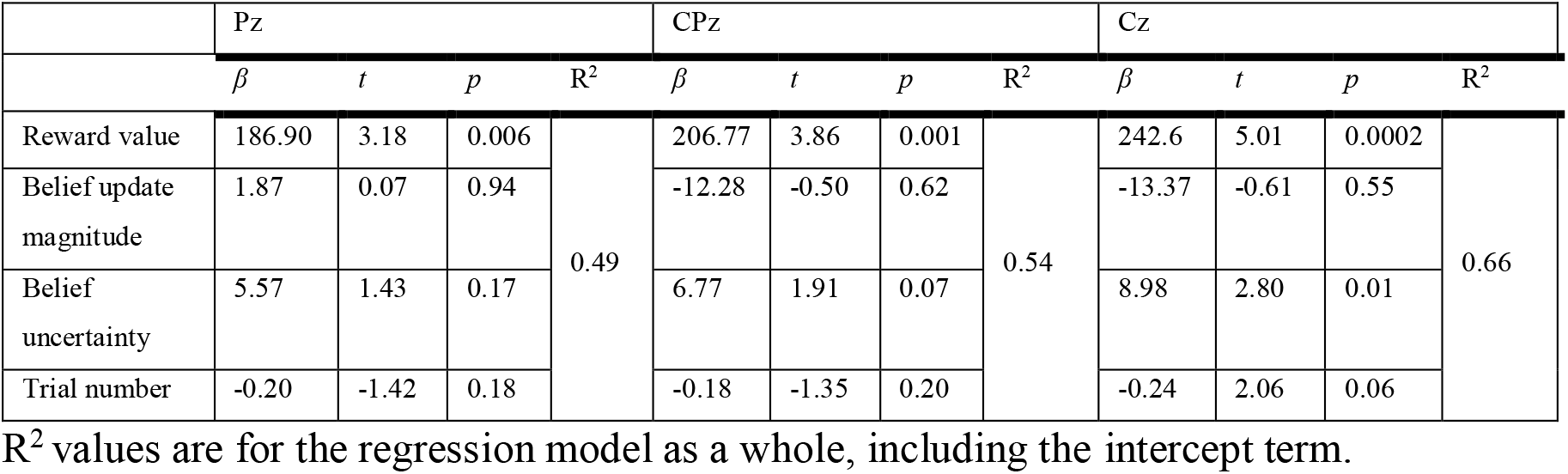
Regression analyses of LPP component in Monetary feedback condition

## 4. Discussion

Within a Bayesian computational modelling framework, the present study set out to identify the effects of motivational factors on the updating of beliefs and on the electrophysiological correlates of belief updating. Concretely, we aimed to differentiate between the effects of monetary reward and the effects of the information content of feedback alone. We quantified these variables with respect to behaviour in a simple reward learning paradigm, and studied the effects of a monetary incentive manipulation on three components of the event-related potential with an established role in learning and/or evaluating the reward value of stimuli: the P3a, FRN, and the LPP (Achtziger et al., 2015; Bennett et al., 2015; Frank et al., 2005; Goldstein et al., 2006; Hajcak et al., 2009; Ito et al., 1998; Jepma et al., 2016; Keil et al., 2002; Polich, 2007; Sato et al., 2005; Yeung & Sanfey, 2004). We found that participants’ choices were more accurate when feedback was delivered in the form of monetary reinforcement than when it was delivered as instructive directives. Belief updating in the reward learning task was also modelled using a Bayesian grid estimator as an additional tool to probe the effect of the feedback incentive manipulation on both behavioural and neural indices, providing model-derived estimates of belief uncertainty (defined as the Shannon entropy of the belief distribution) and belief update magnitude (defined as the mutual information of the feedback) during learning. Model-based estimates of belief uncertainty predicted behavioural performance on the task in both feedback conditions, such that greater belief uncertainty was associated with a larger choice error. Similarly, results revealed that the processing of monetary and instructive feedback could be differentiated by their neural correlates as reflected in the ERP components assessed. On average, amplitudes for the P3a and the LPP were greater given monetary compared to instructive feedback, while the opposite pattern was observed with the FRN.

The event-related potential analyses employed here suggest that, like behaviour, neural encoding of feedback in belief updating differed depending on the presence or absence of monetary reward, even when the net amount of information provided to the decision-maker was equivalent. The P3a has been linked in past research to the process of Bayesian belief updating with a frontocentral topography (Bennett et al., 2015; Kolossa et al., 2015), and it has been proposed that P3a amplitude indexes the magnitude of belief updates, possibly reflecting the deployment of working memory in the revision of prior beliefs (Kopp, 2008). Typically, Bayesian inference involves updating a full belief distribution, a notion also in line with the proposal by Kok (1997) that P3 amplitude may reflect general cognitive effort, since Bayesian belief updating requires a greater expenditure of cognitive resources than other potential mental strategies such as simple heuristics. Interestingly, studies that have identified a centroparietal influence on P3 amplitude tended to rely on paradigms involving infrequent stimuli in order to elicit the P3 such as variations of the oddball task (e.g. Kleih et al., 2010; Mars et al., 2008); however, an effect of belief update on P3a amplitude has also been demonstrated in the same reward learning task as that used here (Bennett et al., 2015). Note that because we implemented two conditions in the present study, with one condition showing smaller belief updates than in the previous study, we could not implement the same analysis as Bennett et al. (2015). Our results, however, clearly demonstrate the effect of monetary reward on the P3a component.

Differential encoding of feedback in the LPP, by contrast, may reflect sustained sensitivity to the reward valence of feedback rather than on belief updates per se. In tasks assessing encoding of affective stimuli, LPP amplitude has been associated with the affective salience of stimuli, such that both positively- and negatively-valenced stimuli elicited larger LPPs than neutral stimuli (Keil et al., 2002; Schupp et al., 2000). Our results showed a differential effect of feedback on the LPP, with overall greater amplitudes for monetary compared to instructive feedback. Multiple linear regression analyses also indicated that belief uncertainty was positively related to LPP amplitude at electrode Cz when other regressors including belief update size and the value of feedback reward were accounted for.

In addition, we observed an overall effect of feedback condition on FRN amplitude, with a larger FRN for instructive compared to monetary feedback. Since we measured the FRN as a peak-to-peak change in amplitude from the immediately preceding positive peak, to say that the FRN component was larger in the instructive condition is equivalent to concluding that the FRN was more negative in the instructive condition than the monetary condition. This effect is therefore in the same direction as that observed for P3a and LPP components, which were measured as mean voltages rather than peak-to-peak voltage changes. This finding is in line with the hypothesis that FRN amplitude reflects a relatively automatic binary evaluation of stimulus valence (Yeung & Sanfey, 2004), and may provide an electrophysiological index of affective components of feedback processing (Keil et al., 2002; Schupp et al., 2000; Wiswede, Münte, Goschke, & Rüsseler, 2009). The differential FRN elicited by monetary and instructive feedback in the present study may be indicative of the greater overall hedonic value of monetary feedback relative to instructive feedback. At two central electrodes, we further found that the model-derived estimate of belief update magnitude was positively related to FRN amplitudes when reward value was taken into account. Such a result suggests a role of the FRN in belief updating *per se* beyond motivational aspects and warrants further research into how the FRN might be linked to integrating stimulus valence into belief updating.

What is the mechanism driving the effect of reward on performance in our task? A key feature of the present reward learning paradigm was that reward (in the monetary condition) was always paired with feedback, while this was not the case in the instructive feedback condition, similar to other reward learning studies (e.g., Bonner, Hastie, Sprinkle, & Young, 2000).

Further, the current task required continuous update of beliefs in an optimally Bayesian manner, instead of simple binary decisions. When feedback is utilised as both an indication of monetary gain and as a learning tool, it is reasonable to infer that participants would have a greater motivational incentive to perform optimally, potentially because the small gains in reward were present during learning. Such an increased motivational drive might have led to an increased investment in cognitive resources, and potentially to either an optimised use of Bayesian strategies, an increase in trials in which a Bayesian strategy was utilised for individual participants, or an increase in the number of individuals who utilised such strategies. In the instructive feedback condition, on the other hand, participants might have felt less motivation to use a computationally demanding Bayesian updating strategy (or less participants might have consistently done so), because they could only rely on intrinsic reward to execute the task correctly, leading to relatively weaker performance. This notion is specifically in line with reinforcement learning theory where individuals, as biological agents, respond to environmental stimuli in ways that will result in the maximisation of reward and minimisation of loss (O’Hara, Hall, van Rijsbergen, & Shadbolt, 2006; Ravindran, 2013). However, we note that there are a number of different ways in which participants’ behaviour may have deviated from Bayes-optimality, and the results of this study do not serve to fully disambiguate between these. For instance, one possibility is that participants in the instructive condition were more likely to adopt an entirely non-Bayesian heuristic strategy (such as Win-Stay-Lose-Shift); another is that they performed a form of probabilistic belief revision that underweighted the uncertainty of the prior distribution; finally, still another possibility is that Bayesian belief revision was employed in both conditions, but that the precision of updating was lower in the instructive feedback conditions. Future studies could further investigate these hypotheses. In particular, investigating the instructive/monetary feedback manipulation in a larger sample size would have more power to detect a difference between conditions in the *σ* parameter, which was not statistically significant in the present study.

From a psychological and neural standpoint, there are two distinct sub-processes within the task that might have been affected by the feedback manipulation. The first is the learning process: monetary feedback may have improved learning, leading to participants’ having more accurate representations of the reward structure of the task. The second is the choice process: participants may have translated this learned representation into choices more accurately in the monetary feedback condition, since the incentives for them to do so were stronger. (In the framework of Bayesian decision theory, this distinction between the learning process and the choice process corresponds to a distinction between belief updating and the cost function.) For the choice data that we collected, it is difficult to distinguish these two explanations, since changes in either process will produce indistinguishable changes in behaviour in our task. This identifiability concern motivated our decision to fit choice data using a computational model with the same σ parameter for both the inference process and the choice process. This point notwithstanding, our neural results shed some light on the distinction between motivational effects on learning versus motivational effects on choice. Since our ERP data showed that the neural representation of feedback differed between feedback conditions at the point of feedback presentation, our results are, at least in part, consistent with an effect of motivational state on learning. Of course, this finding requires further investigation, and does not preclude the possibility that motivational state influences choice in other settings.

In turn, there are several underlying neural mechanisms that might be responsible for this effect on learning. One possibility is that the monetary feedback enhanced participants’ attention to the task, and that this greater engagement manifested in increased neural gain for feedback in the monetary condition (potentially suggesting a role for norepinephrine; see Eldar, Cohen & Niv, 2013; Jepma et al., 2016). Another is that monetary feedback affected the deployment of visual working memory resources, such that the chosen contrast was represented with greater precision in the monetary feedback condition (Bays, Catalao & Husain, 2009), thereby leading to more precise belief updating.

More broadly, our results have bearing on the hypothesis that Bayesian inference represents a unifying principle of neural computation (the ‘Bayesian brain’ hypothesis; Knill & Pouget, 2004). This hypothesis has been applied successfully to domains including sensory coding and motor planning (Körding & Wolpert, 2004; Yuille & Kersten, 2006). In computational terms, neural processing costs comprise both the expense of computing action policies, and the difficulty of learning (computational complexity versus sample complexity). This situates Bayesian models of cognition within an ecologically valid framework in which inference is constrained by the cognitive resource limitations of the brain. However, one issue with applying Bayesian inference to higher-level judgement and decision-making is that Bayesian inference is resource-intensive, and therefore computationally intractable for many real-world tasks (see e.g., Bossaerts & Murawski, 2017; Payzan-LeNestour & Bossaerts, 2011). Indeed, much evidence suggests that in many decision settings, humans fail to employ optimal Bayesian strategies (e.g., Cassey, Hawkins, Donkin, & Brown, 2016; Gigerenzer & Goldstein, 1996). Moreover, even in cases where Bayesian inference is tenable, many individuals instead appear to rely on heuristic strategies (e.g., Bennett et al., 2015; Steyvers, Lee, & Wagenmakers, 2009). Although our results provide evidence for Bayesian inference in belief updating in general, within the context of the present experiment, these previous studies support the idea that judgement and decision-making are not always performed using optimal Bayesian strategies. Our results further suggest that the reward contingencies of the environment might have a strong influence on whether or not beliefs are updated using optimal Bayesian strategies, even within the same individuals.

Our results are also broadly in line with the recently proposed economic value of control framework (Shenhav et al., 2013; 2017). This framework proposes that cognitive resources are allocated to tasks according to the expected value gained from resource use, which is given by the difference between expected reward and expected costs of resource use. The latter arise from both metabolic as well as opportunity costs. In our task, we manipulated reward while cognitive resource requirements were kept constant across conditions. Thus, expected reward of resource use was higher in the monetary feedback condition while expected costs were the same. The finding that participants behaved more optimally in the monetary feedback condition could thus be interpreted as a result of participants allocating more cognitive resources to the task in the monetary feedback condition due to higher expected value, in turn using a more computationally-intensive strategy than in the instructive feedback condition. Our finding of a higher P3a amplitude, which has previously been associated with cognitive effort, in the monetary feedback condition further supports this interpretation (Shenhav et al., 2017).

Finally, the results of this study touch on an ongoing debate in behavioural economics and educational psychology concerning the distinction between intrinsic and extrinsic motivation for performance. A classical finding in this respect is that, if an individual is already intrinsically motivated to perform a task, the introduction of extrinsic motivators (such as monetary performance incentives) can undermine performance by causing a decrease in intrinsic performance motivation (Deci, 1971). More recent findings have refined this notion by suggesting that intrinsic and extrinsic motivation account for non-overlapping variance in performance (Cerasoli, Nicklin & Ford, 2014), or that there is a non-linear interaction between intrinsic and extrinsic motivation in determining performance (Lin, McKeachie & Kim, 2003). By showing that monetary incentives improved perceptual learning, our results suggest that extrinsic motivation can lead to improved performance in simple cognitive tasks. However, we interpret the applications of this result only with great caution, given the dissimilarity between our reward learning task and applications such as the classroom or workplace. One potential link is the literature on ‘gamification’ of classroom and workplace tasks, in which extrinsic motivators take the form of computer-game elements such as badges, points, or levels (Buckley & Doyle, 2014; Dicheva, Dichev, Agre & Angelova, 2015). Our results suggest a candidate neuropsychological process that could underlie these effects.

In summary, the present study provides evidence that motivational factors, such as the presence of monetary reward in performance feedback, can increase learning rates in tasks demanding continual belief updating. This enhancement of performance was independent of the information content of the feedback itself. The presence or absence of monetary reward was reflected by components of the event-related potential previously linked to motivation, feedback processing and stimulus salience, the P3a, FRN and LPP, respectively. Overall, our results suggest that motivational state may critically affect the use of Bayesian inference in belief updating.

## Acknowledgements

We thank Daniel Feuerriegel and Maja Brydevall for help with ERP analysis. This work was supported by a Strategic Initiatives Fund grant from the Faculty of Business and Economics at The University of Melbourne to C.M. and S.B., and an Australian Research Council Discovery Early Career Researcher Award (DE 140100350) to S.B.

